## Supplementary Information for "Sequence-Dependent Material Properties of Biomolecular Condensates and their Relation to Dilute Phase Conformations"

**Table S1.** Amino acid codes of E−K, LAF1, and DDX4 sequence variants that were used in this study.

| **E−K** | **Sequence** |
|---|---|
| nSCD = 0.000 | EKEKEKEKEKEKEKEKEKEKEKEKEKEKEKEKEKEKEKEKEKEKEKEK<br>EKEK |
| nSCD = 0.017 | KEKKKEKEEEKKEKEKKEEKEKEEKKEEKKEEEKKKEEEKKEKKKE<br>EEEK |
| nSCD = 0.033 | KEKKKEKEEEKEEEKKEEKEEEKEEKKKKEEEKKKEKEEEEKEKKK<br>EKEK |
| nSCD = 0.059 | KKEEKEKEEEKEEEEEEKKKKEEKEKEKKKEKKEEEKKKEEKKKKE<br>EEKK |
| nSCD = 0.067 | KKEEKEKKEEEEEKEEEKEEKKKEEEEKKKKKKKKEKKEKEEEEKEK<br>KEEK |
| nSCD = 0.200 | EEKKEKEEEEEEKEKEEEEEEKEEKEKKKKKKKKEKEKKKEKKKKEK<br>EEKK |
| nSCD = 0.333 | EEEKEKEKEKEEEEEEEEEEEKEEKEKEKKKKKKEEKKKEKKKKKE<br>KKKK |
| nSCD = 0.400 | EKEKEEEEEEKEEEEEEEEEKEEEKKEKEKKKKKKEEEKEKKKKKKK<br>KKKK |
| nSCD = 0.468 | EEEEEEEEEEKEEEEEKEEKKKEKKEKEEEKKKKEEKEKKKKKKKK<br>KKKK |
| nSCD = 0.499 | EEEEEEEEEEKEEEEEEEEEKKKEKKKKEEEKKKKEKKEKKEKKKKK<br>KKKK |
| nSCD = 0.601 | EEEEEEEEEEEEEEEEEKEEKKEKEEKKKEKKKEKEKKKKEKKKKKK<br>KKKK |
| nSCD = 0.667 | EEEEEEEEEEEEEEEEEKEEEKEKEKEKEKKKKKKKKEKKKKKKKKE<br>KKKK |
| nSCD = 0.734 | EEEEEEEEEEEEEEEEKEEEEEKKKEEEEKKKKKKKKKKEKKKKKKK<br>KKKK |

| | |
|---|---|
| nSCD = 0.902 | EEEEEEEEEEEEEEEEEEEEKEKEKEKEKEKKKKKKKKKKKKKKKK<br>KKKK |
| nSCD = 1.000 | EEEEEEEEEEEEEEEEEEEEEEEEEKKKKKKKKKKKKKKKKKKKKK<br>KKKK |
| **LAF1** | **Sequence** |
| nSCD = 0.003 | MGGSGGRGSGRNGDYYGSGPGYRAQDYGHGDSRRGGANSSGGNG<br>GNPGRAEGREGGGGANGYYSGGGANQASGGNRNGGADGGGGGDR<br>GGGNNGNRGDAGYREGNRRGDNNGNFGDNNSDRGLRGASDRDGN<br>RNYLGSRDRDVNGGGDYNYRNGGYRNRGDDGRSGDR |
| nSCD = 0.010<br>**Wild-type** | MESNQSNNGGSGNAALNRGGRYVPPHLRGGDGGAAAAASAGGDD<br>RRGGAGGGGYRRGGGNSGGGGGGGYDRGYNDNRDDRDNRGGSG<br>GYGRDRNYEDRGYNGGGGGGGNRGYNNNRGGGGGGYNRQDRGD<br>GGSSNFSRGGYNNRDEGSDNRGSGRSYNNDRRDNGGDG |
| nSCD = 0.039 | MRRGGYGGDGGLGNGHNDERRNARRGQNNSGFGYDAGRRSRGRY<br>GAGGGDRNGRNDSNGLGGNSGDNGGGRNGANRGYRGDDNNARGS<br>NVGRYSDDGARREGSSSGGGGYGNNGGGNGDGPGANGRDAYRGN<br>DGGPYQRSGYAGSNGGRDGDNSDYYDGGNRGEGGGG |
| nSCD = 0.131 | MGGDRAGYNNGDRLNRNGFRRLSGPNGAGGGNSDRRYYRNARRG<br>GGRGGRAGGYGYNGGRRRAQGSRGEGGGGGGSYDNGNRGEGSGR<br>GRNNDGGGDDNARNNDSHRDPGRVGNAGYGGGGGNGNGSYYGG<br>DNNDAGGSGSADGGGSNGDYDERGDSNGRDGNGQYDS |
| nSCD = 0.206 | MNNSGDNDRGSGNYGLRNSFGDDGYGDNGNDEGNSGYRNRGLGG<br>DRADEYGNSGGNGDNEAAPNASDRDDAHYYDSDDYDDGGGGRGS<br>GGAGGGGARGPGSNRAGRYGGGGRRGRGRGNGYNGNRSQRRRGG<br>GRGRGNRGYRVGNGNGQSGGRNSRGGGGGNGGANYG |
| nSCD = 0.263 | MDGGDDDGGQGGAGENLNDYYNGSSGGPGNSVAENSGGNGNDDG<br>GGDNRNNRYYSGGGDGRYGGGGQGANGGDDNAGDGYDGDNFGN<br>GNGRGYNNGSYGADASEGDRGSYRSSGRRRRRGSGGGGNYPGGGS<br>RRLNRNRGARGGAGGNRRRNARRDGNGYGGRRGRGH |
| nSCD = 0.394 | MESNQSNNGGSGNAALNDGGDYVPPHLDGGDGGAAAAASAGGDE<br>DDGGAGGGGYDDGGGNSGGGGGGGYDEGYNDNDDDDNDGGSGG |

| | |
|---|---|
| | YGDRRNYRRRGYNGGGGGGGNRGYNNNRGGGGGGYNRQRRGRG<br>GSSNFSRGGYNNRRRGSRNRGSGRSYNNRRRRNGGRGR |
| **DDX4** | **Sequence** |
| nSCD = 0.002 | MDPTESFCANGGGAGSNKMGDGGNRATSDGLKNVFSMSNEADGRS<br>SDRSGNSNSGDPKDKGGETSRESRNGSGGFERGREPGNSGRNIGSSTI<br>EERLSRISGSDGTPYLDGSYSRERVSADHGGNMFYCFPSDNKGGSGL<br>RPNANEFNRERTSQVMDRPNGEGFFRSFYRHPVGGMESTYWGQSGS<br>GPERNGFGSDDGWRLSFRKDEWRCSDKDNGFNEFRKSGNGGERCT<br>GEGDF |
| nSCD = 0.010 | MGRDRLRGDNDLPKGYEGFENEHFSKSNTKGTRGPVNNEGDRGFN<br>DGSYNGIGSRGQFGDGSCEPGGNMKMGVTGGGCSISEEGPSEENNR<br>RNRWRQFGNDSGDGWVPGPACHRGISGFFRKDTSGKSDCDERTSAR<br>SREEGDSGLDSSYWGEGSFRSASRRRNRGDSNELDSGSEFYGGNENP<br>GMRSDAFTGSGEPFSPDSSKMNFTSSAYFFANDVKELGSNTSRGRNG<br>GGRMS |
| nSCD = 0.021<br>**Wild-type** | MGDEDWEAEINPHMSSYVPIFEKDRYSGENGDNFNRTPASSSEMDD<br>GPSRRDHFMKSGFASGRNFGNRDAGECNKRDNTSTMGGFGVGKSF<br>GNRGFSNSRFEDGDSSGFWRESSNDCEDNPTRNRGFSKRGGYRDGN<br>NSEASGPYRRGGRGSFRGCRGGFGLGSPNNDLDPDECMQRTGGLFG<br>SRRPVLSGTGNGDTSQSRSGSGSERGGYKGLNEEVITGSGKNSWKSE<br>AEGGES |
| nSCD = 0.095 | MGDPDDDEEEKGSNSTGGEQDGELGQDVSANDLGGSRSTSEDEFSR<br>EYNEGPSYGYSTGRFKDWRFTEFSFREFTFMNGPGNKGEIDGRWHIF<br>DNAGCSDSGPRGDITESGGSSSSNNKRAGGVKNRNDGEGDSMPGGR<br>GTRNGRHRGGSPNRRPFVAGSNSDRNNLSCNGSLEWYGGEGPCRSP<br>RMGGGAGNMSGGYRKGDERMSLVNGATGKFNRDERRRSFSSCFFN<br>SSFKSS |
| nSCD = 0.194 | MRNESEGEYCCEADDFVGNPSDNSYTRASTGSDDDRGFYFSFEAGG<br>SYDEFSSEGDEWDNNGSGGGDNFNESDFRGDNETGDPERGDNDGIE<br>KHGDPGISPLGRSPGMSRTTDTRGSMLGGSFEVGSGFLFTGESGSAS<br>MGWRGNSRFKFCRDMLQASKRINNNKVSGGRRSGESWCSGGREGT |

| | |
|---|---|
| | GSSMRGRGPKFEGNRGRSPNNKAQGKRRRPFGGKHNRYNSERNGSL<br>NVSSGP |
| nSCD = 0.284 | MNSEEYCGCEADDDFVGNPSNSYTRASTGSRDDEDGFFSYDEEEEEE<br>EFSSGDWDNNGSGGGEEDNFNSDFIKHRGDNETGDGSPGFGNGYFS<br>GDDPGISPLGSPGMSERRRRTTDDDTGGGSFVAGGLFTGSGSASMG<br>WSMFKRRFLCRRRKMLQASGNSINVSSNNKVSGGSGESWCSGGRRG<br>TGSSMKGGPFEGNGSPNNKAQGKRRRRRPFGGKHNRRRRYNSEN<br>GSLNGP |
| nSCD = 0.478 | MNESEEGYCACEDDEFVGNPSDNSEEYTASTGSDDDEEEEEEDEGFY<br>FSFAGGSYFSSGDWDNNGSGGGDNFNSDFGDNETGDPEGDNDGIEK<br>HGDDPGISPLGSPGMSRTTTRRGSMLGGSFEVGSGFLFTGSGSASMG<br>WRRRRGNSFKFGSWCSCDMLQASINNVGGNKNKVSSLRRRRRSGG<br>GTGSSMGKRRGPFGGNSPNNKAQGRRRRRRRRRPFGGKHNYNSRR<br>NGSSGP |

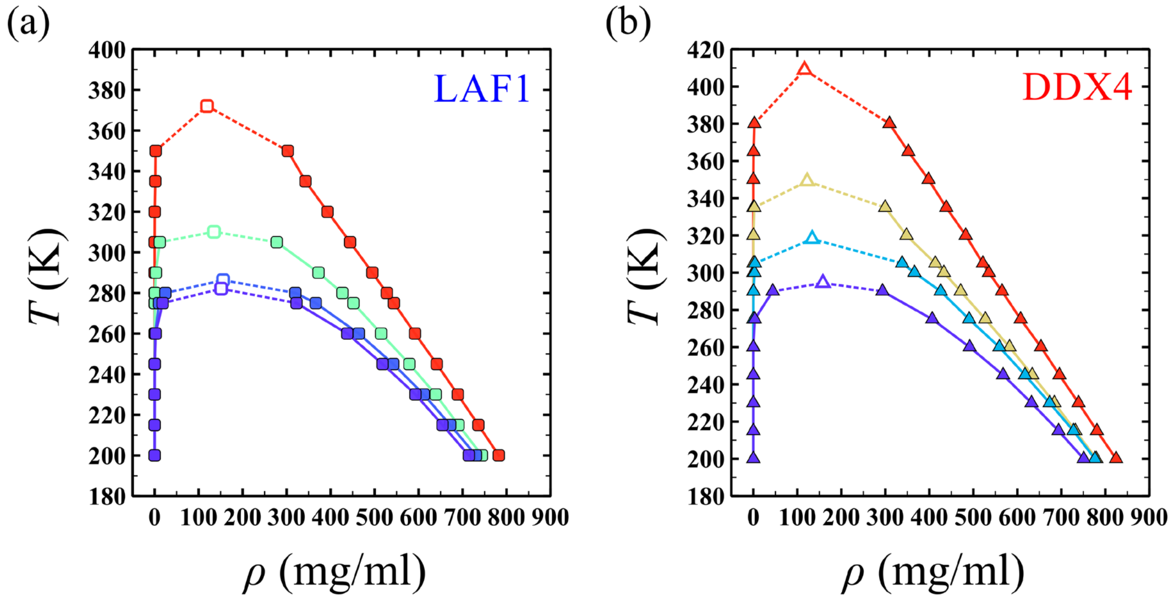


**Fig. S1.** Phase diagrams for select (a) LAF1 (nSCD $= 0.003, 0.010, 0.131,$ and $0.394$) and (b) DDX4 (nSCD $= 0.002, 0.021, 0.194,$ and $0.478$) sequence variants. The open symbols correspond to the critical temperature and critical density of the sequence variants. The line color, ranging from blue to red, indicates increasing nSCD. Lines are guide to the eye only.

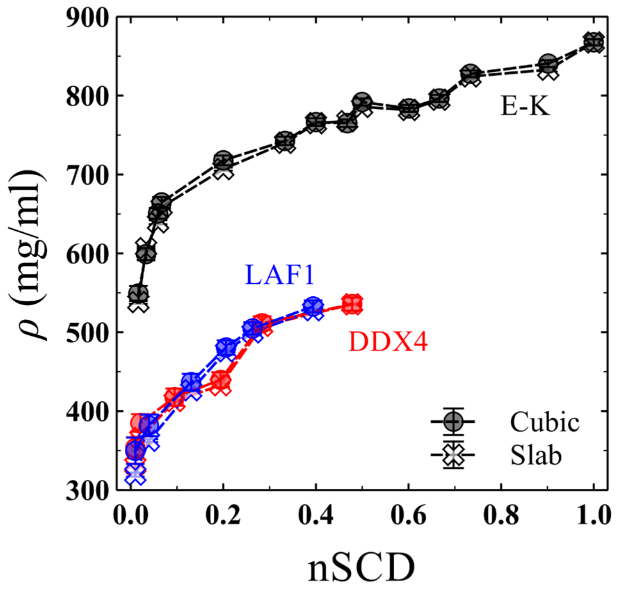


**Fig. S2.** Comparison between the dense phase concentration $\rho$ attained by the E−K, LAF1, and DDX4 sequence variants in cubic and slab geometries.

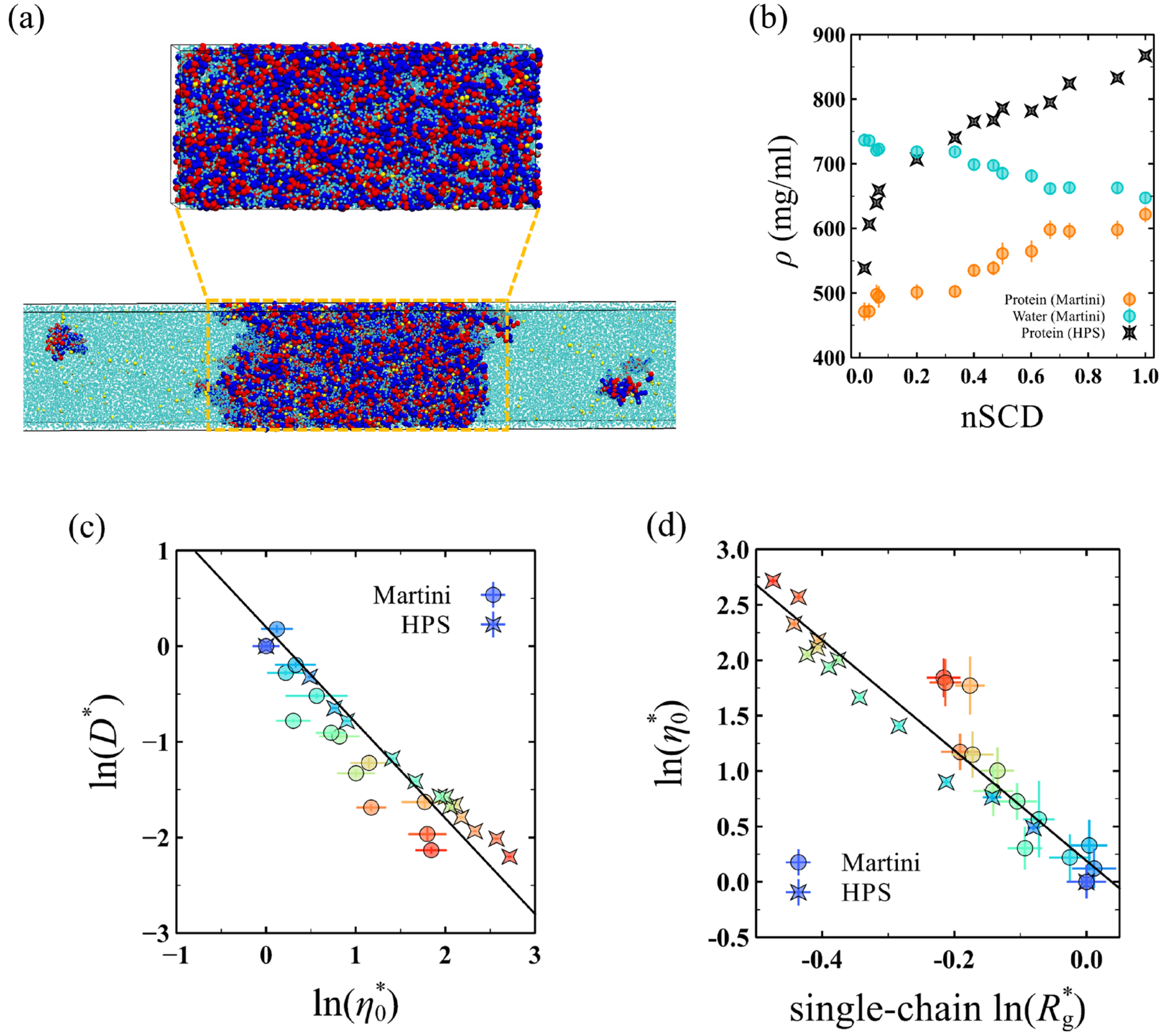


**Fig. S3.** (a) Dense phase snapshot of Martini set-up for one of the E−K sequences in the presence of water (cyan) and ions (yellow) in a cuboid simulation box, which was cut from a slab geometry, for characterizing its dense phase diffusive and rheological properties. (b) Comparison of the dense phase concentration $\rho$ for protein and water as a function of nSCD in the Martini model with the protein's concentration in the HPS model for the E−K sequences. (c) Inversely proportional correlation between normalized diffusion coefficient $D^*$ and normalized zero-shear viscosity $\eta_0^*$ from the Martini and HPS models for the E−K sequences, with the solid line corresponding to the Stokes-Einstein type relation $D^* = 1/\eta_0^*$. (d) Correlation between $\eta_0^*$ and normalized single-chain $R_g^*$ from the Martini and HPS models for the E−K sequences. The solid line corresponds to the linear fit $\ln(\eta_0^*) = -4.98\ln(R_g^*) + 0.19$ with correlation coefficient $R^2 = 0.90$. The values of $D^*$, $\eta_0^*$, and $R_g^*$ are obtained after normalizing $D$, $\eta_0$, and $R_g$ by those computed for the reference E−K sequence with nSCD = 0.017 in the HPS and Martini models. The symbol color, ranging from blue to red in (c) and (d), indicates increasing nSCD.

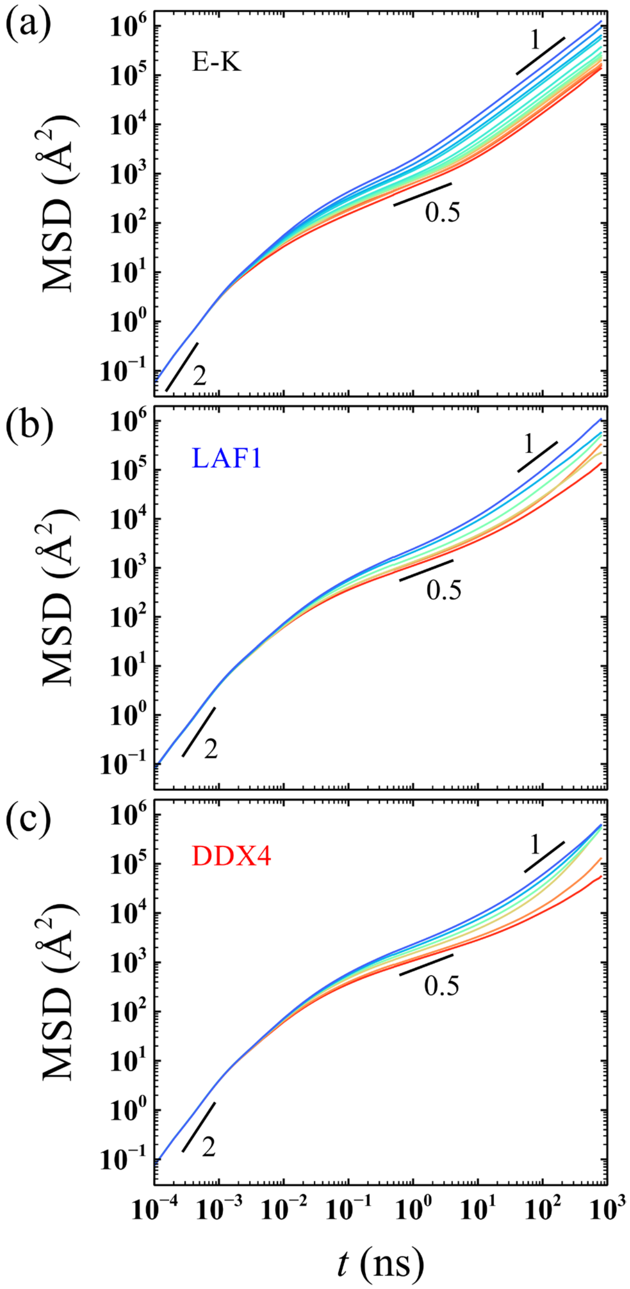


**Fig. S4.** Mean square displacement MSD($t$) of the residues of a chain in the dense phase for the (a) E−K, (b) LAF1, and (c) DDX4 sequences. To avoid chain end effects, 20 residues on either end of a chain have not been considered for computing the MSD. The slope values 2, 0.5, and 1 indicate ballistic, sub-diffusive, and diffusive regimes, respectively. The line color, ranging from blue to red, indicates increasing nSCD.

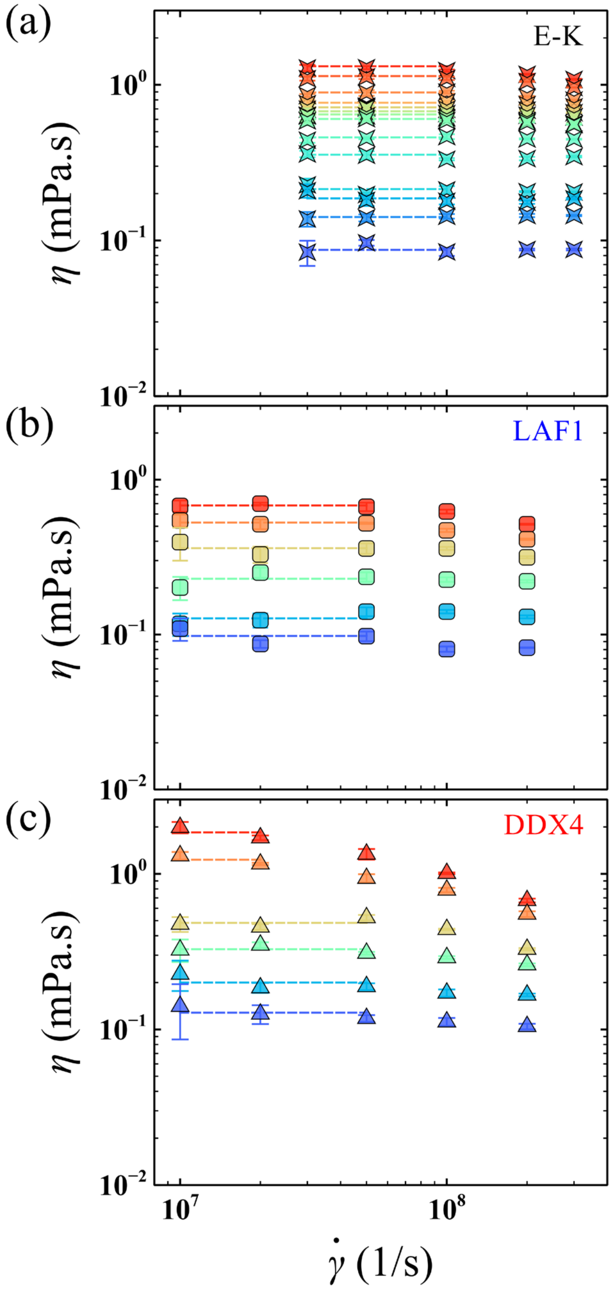


**Fig. S5.** Shear viscosity $\eta$ as a function of shear rate $\dot{\gamma}$ for the (a) E−K, (b) LAF1, and (c) DDX4 sequences in the dense phase. The dashed lines indicate the zero-shear viscosity $\eta_0$ of each sequence, which is obtained as an average of the near-constant $\eta$ values from the Newtonian plateau observed at low shear rates. The symbol and line colors, ranging from blue to red, indicate increasing nSCD.

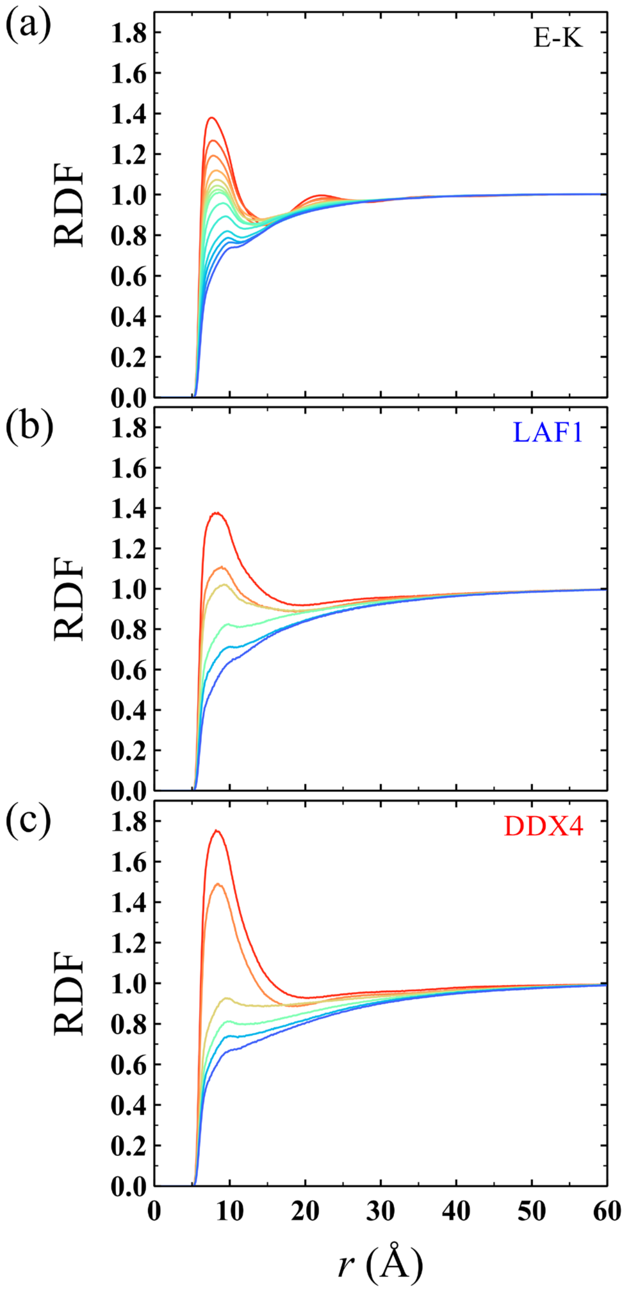


**Fig. S6.** Radial distribution function (RDF) as a function of distance $r$ between the interchain oppositely charged residues for the (a) E−K, (b) LAF1, and (c) DDX4 sequences in the dense phase. The line color, ranging from blue to red, indicates increasing nSCD.

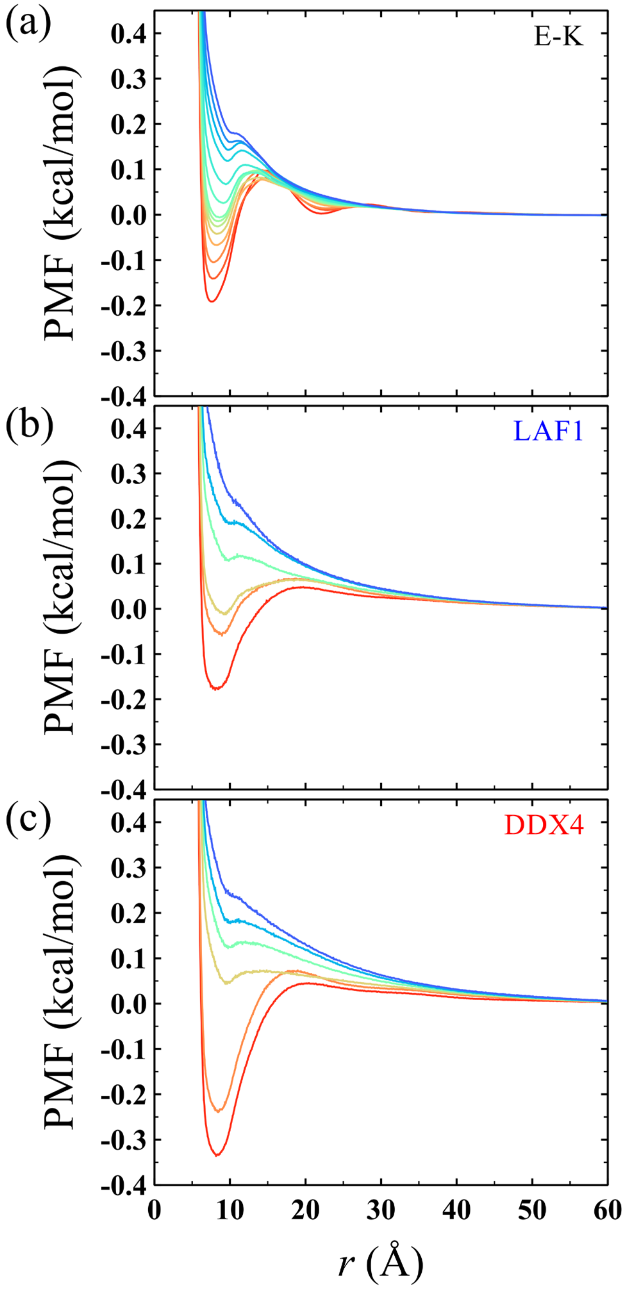


**Fig. S7.** Potential of mean force (PMF) as a function of distance $r$ between the interchain oppositely charged residues for the (a) E−K, (b) LAF1, and (c) DDX4 sequences in the dense phase. The line color, ranging from blue to red, indicates increasing nSCD.

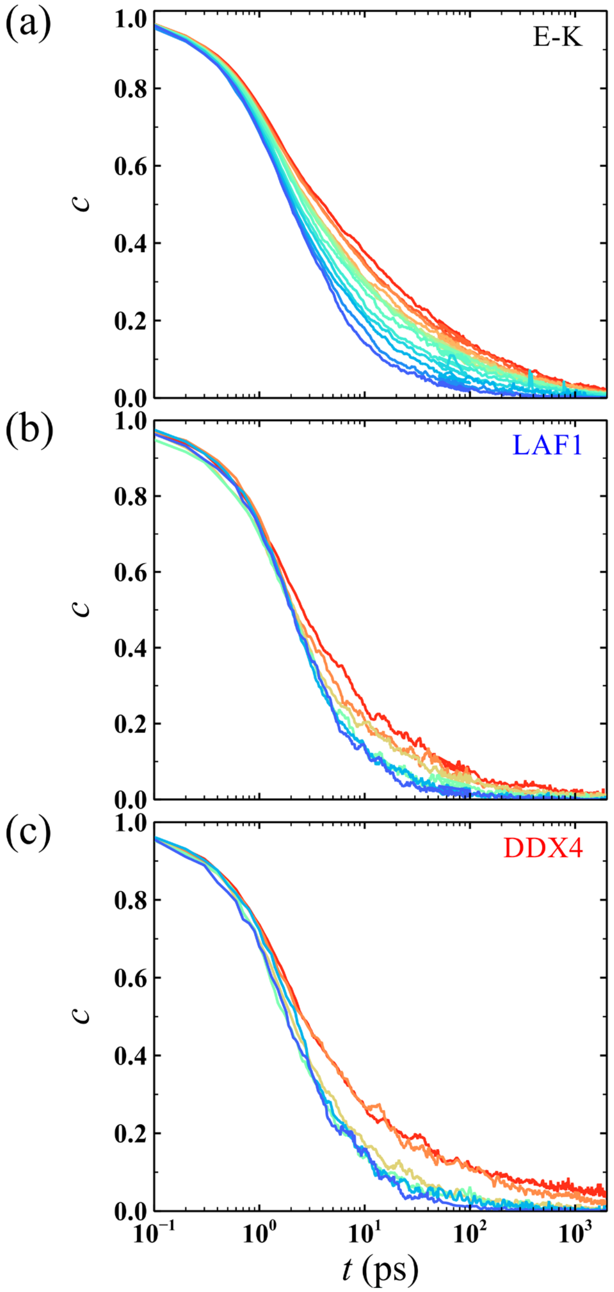


**Fig. S8.** Intermittent contact time autocorrelation $c(t)$ between the interchain oppositely charged residues for the (a) E−K, (b) LAF1, and (c) DDX4 sequences in the dense phase. The line color, ranging from blue to red, indicates increasing nSCD.

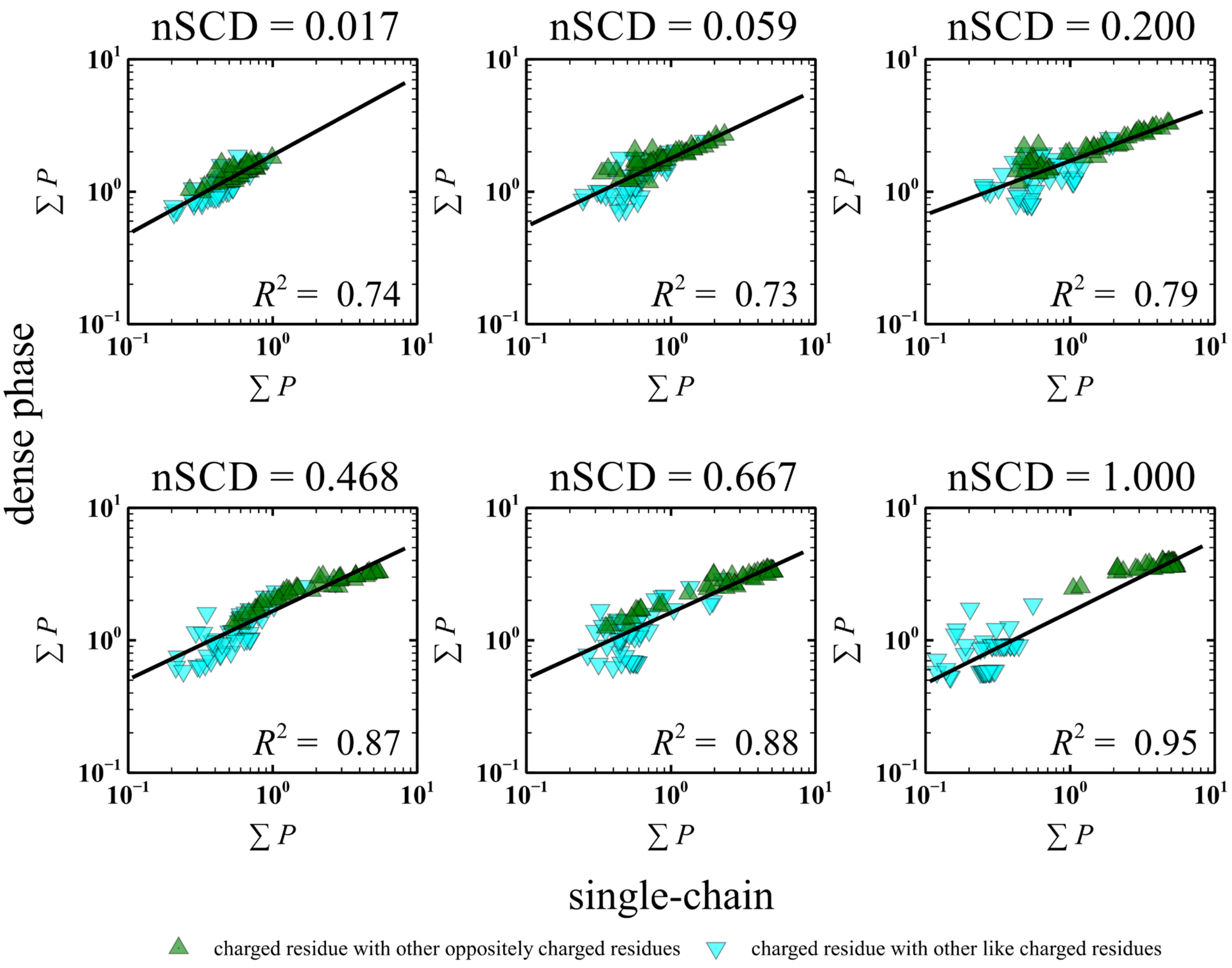


**Fig. S9.** Correlation between the summed-up contact probabilities $\sum P$ of a residue $i$ with all other oppositely charged residues or like charged residues for the chains in the dense phase and for a single-chain of select E−K sequences. For the single-chain $\sum P$, contributions from directly bonded residues and residues that are separated by two or three bonds are excluded as their contacts are primarily established by the chain connectivity.

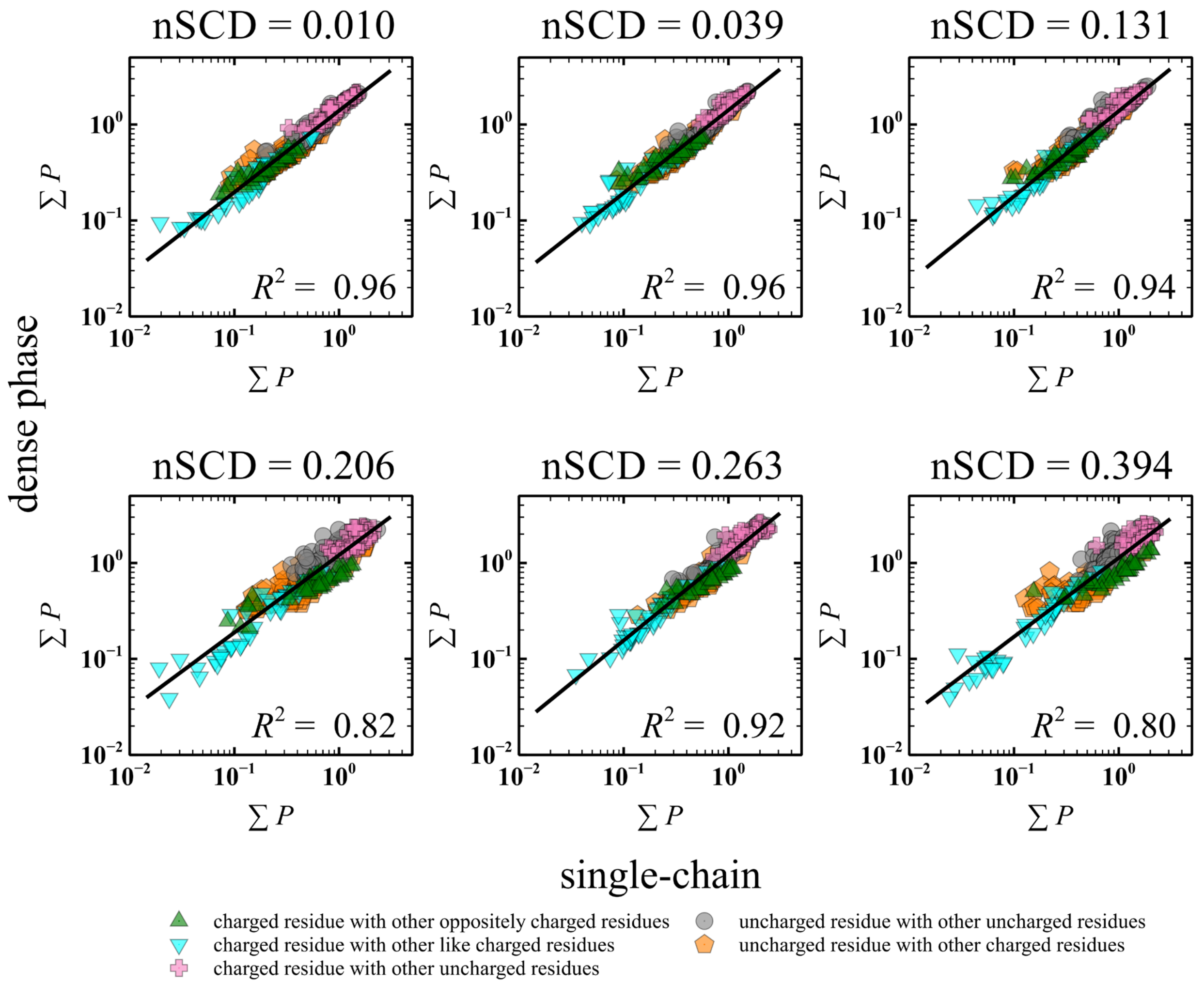


**Fig. S10.** Correlation between the summed-up contact probabilities $\sum P$ of a residue $i$ with all other oppositely charged residues or like charged residues or uncharged residues for the chains in the dense phase and for a single-chain of select LAF1 sequences. For the single-chain $\sum P$, contributions from directly bonded residues and residues that are separated by two or three bonds are excluded as their contacts are primarily established by the chain connectivity.

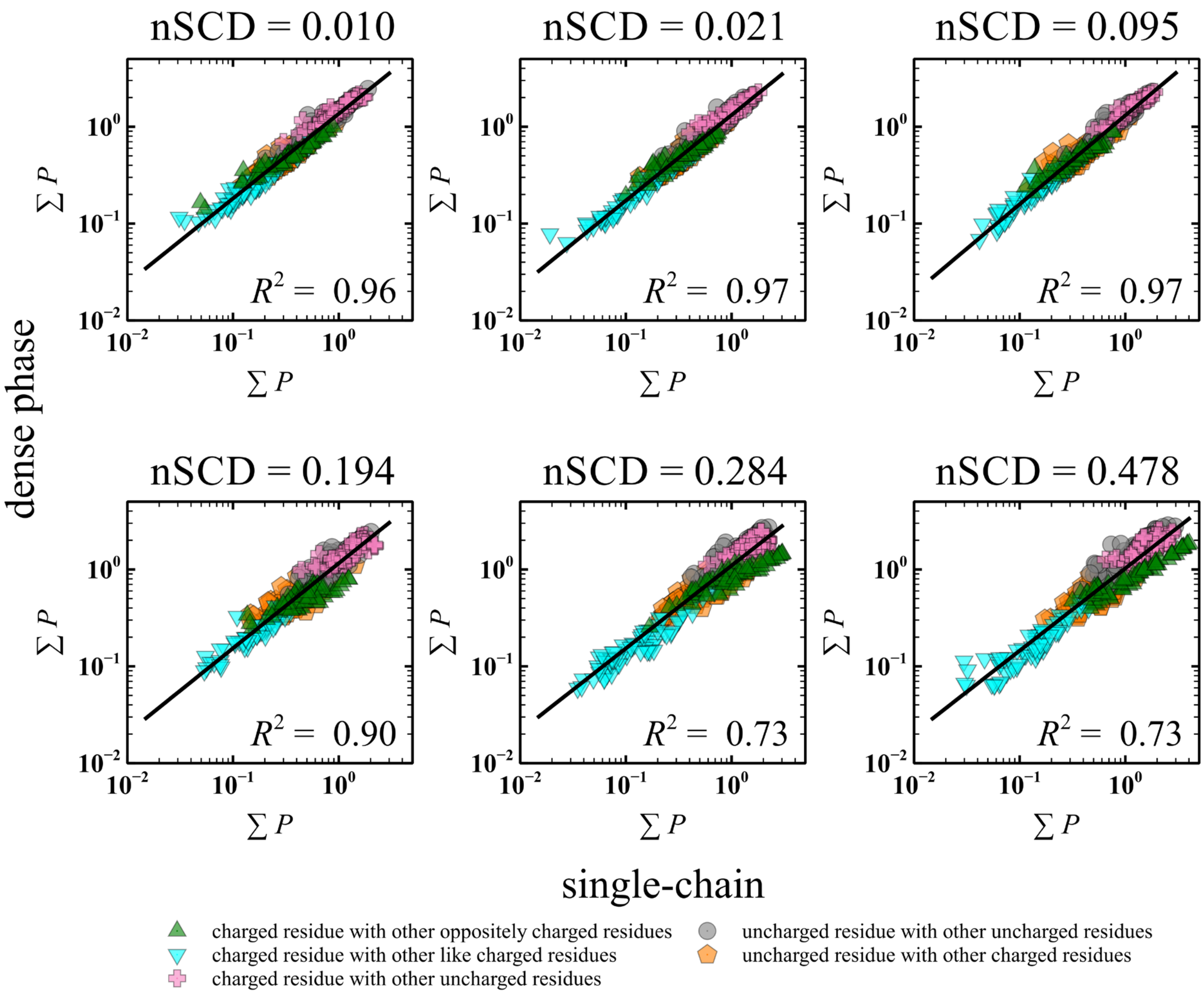


**Fig. S11.** Correlation between the summed-up contact probabilities $\sum P$ of a residue $i$ with all other oppositely charged residues or like charged residues or uncharged residues for the chains in the dense phase and for a single-chain of select DDX4 sequences. For the single-chain $\sum P$, contributions from directly bonded residues and residues that are separated by two or three bonds are excluded as their contacts are primarily established by the chain connectivity.

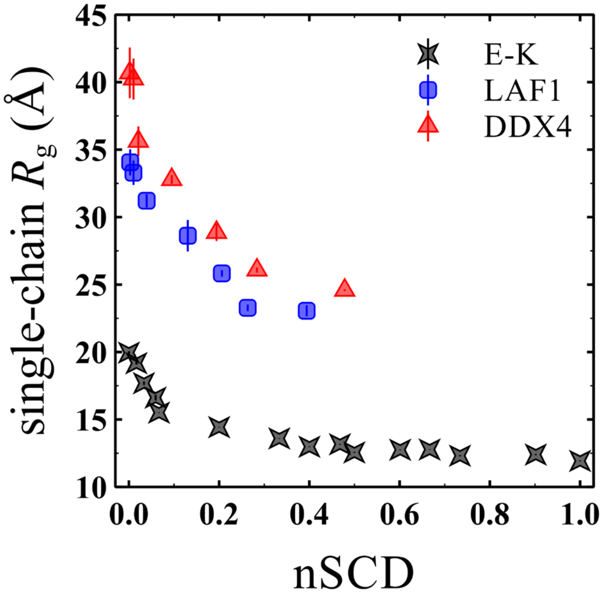


**Fig. S12.** Radius of gyration $R_g$ as a function of nSCD for the single-chain E−K, LAF1, and DDX4 sequences.

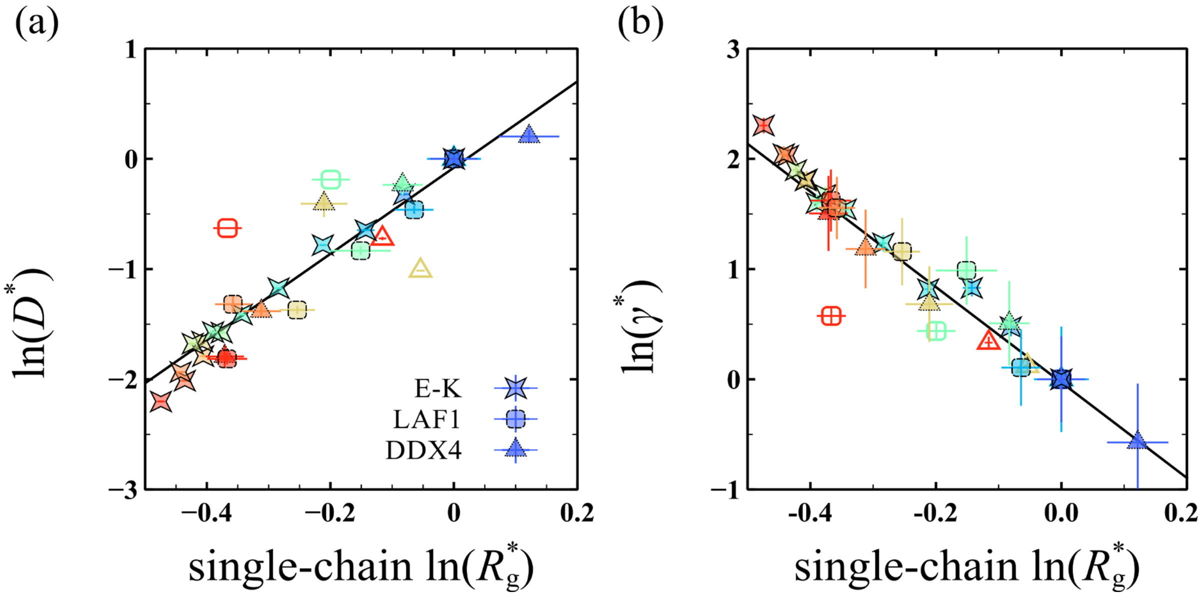


**Fig. S13.** Correlation (a) between diffusivity $D^*$ and single-chain $R_g^*$, and (b) between surface tension $\gamma^*$ and single-chain $R_g^*$ for all sequences. The symbol color, ranging from blue to red, indicates increasing nSCD. The values of $D^*$, $\gamma^*$, and $R_g^*$ are obtained after normalizing $D$, $\gamma$, and $R_g$ by those computed for the same reference sequences used in **Figs. 2 and 3**. The closed and open symbols correspond to the values obtained based on Urry's hydropathy scale and Kapcha-Rossky's hydropathy scale, respectively. The solid lines correspond to the linear fits $\ln(D^*) = 3.91\ln(R_g^*) - 0.08$ with correlation coefficient $R^2 = 0.84$, and $\ln(\gamma^*) = -4.33\ln(R_g^*) - 0.03$ with $R^2 = 0.91$, respectively.

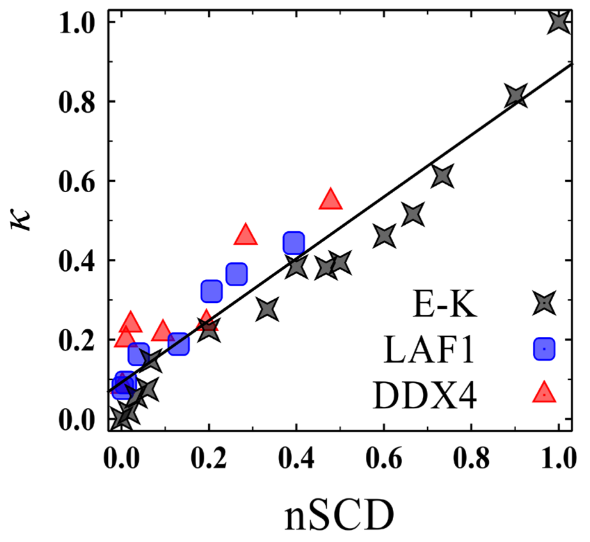


**Fig. S14.** Correlation between nSCD parameter used in this work and $\kappa$ parameter introduced by Das and Pappu.[2] The solid line corresponds to a linear fit with correlation coefficient $R^2 = 0.90$.

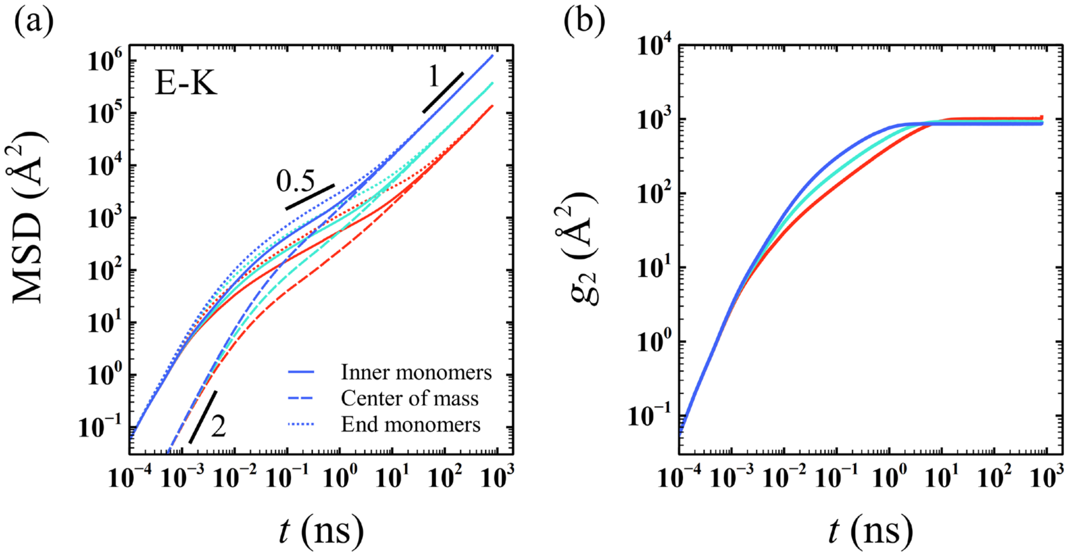


**Fig. S15.** (a) Mean square displacement MSD($t$) of the inner residues of a chain (solid lines), the center of mass of a chain (dashed lines), and the end monomers of a chain (dotted lines) in the dense phase for select E−K sequences (nSCD = 0.017, 0.200, and 1.000). The slope values 2, 0.5, and 1 indicate ballistic, sub-diffusive, and diffusive regimes, respectively. Inner residues correspond to those in the middle of a chain after excluding 20 residues on either end of the sequence. (b) MSD of the inner residues of a chain relative to the center of mass $g_2(t)$ in the dense phase for select E−K sequences. The line color, ranging from blue to red, indicates increasing nSCD.

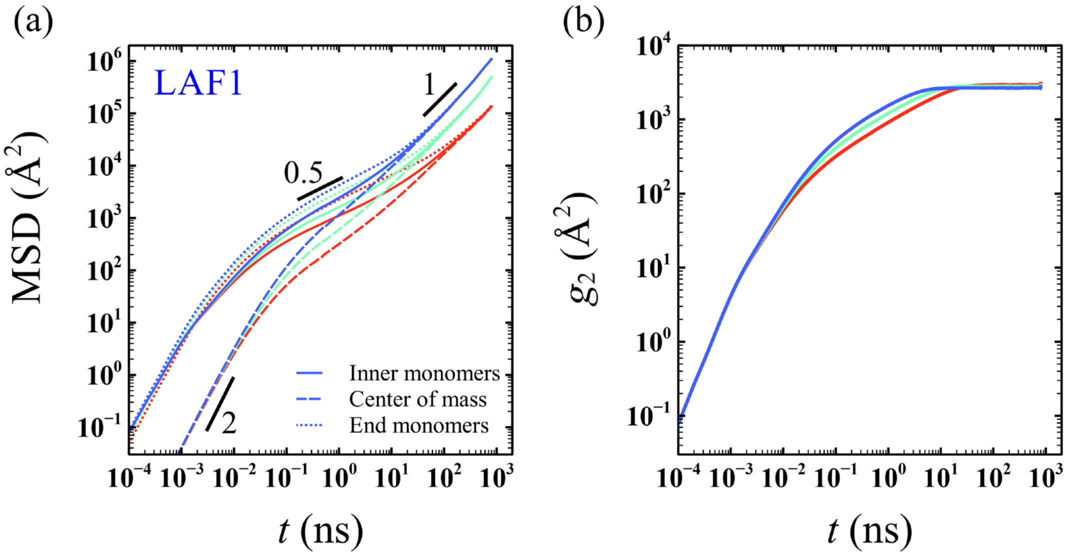


**Fig. S16.** (a) Mean square displacement MSD($t$) of the inner residues of a chain (solid lines), the center of mass of a chain (dashed lines), and the end monomers of a chain (dotted lines) in the dense phase for select LAF1 sequences (nSCD $= 0.010, 0.131$, and $0.394$). The slope values 2, 0.5, and 1 indicate ballistic, sub-diffusive, and diffusive regimes, respectively. Inner residues correspond to those in the middle of a chain after excluding 20 residues on either end of the sequence. (b) MSD of the inner residues of a chain relative to the center of mass $g_2(t)$ in the dense phase for select E−K sequences. The line color, ranging from blue to red, indicates increasing nSCD.

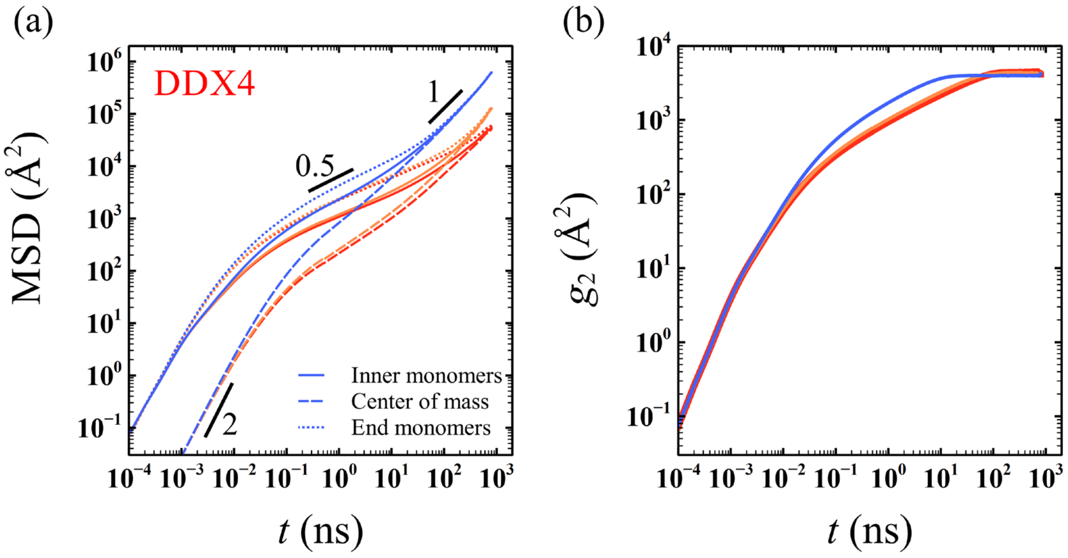


**Fig. S17.** (a) Mean square displacement MSD($t$) of the inner residues of a chain (solid lines), the center of mass of a chain (dashed lines), and the end monomers of a chain (dotted lines) in the dense phase for select DDX4 sequences (nSCD $= 0.010, 0.284$, and $0.478$). The slope values 2, 0.5, and 1 indicate ballistic, sub-diffusive, and diffusive regimes, respectively. Inner residues correspond to those in the middle of a chain after excluding 20 residues on either end of the sequence. (b) MSD of the inner residues of a chain relative to the center of mass $g_2(t)$ in the dense phase for select E−K sequences. The line color, ranging from blue to red, indicates increasing nSCD.

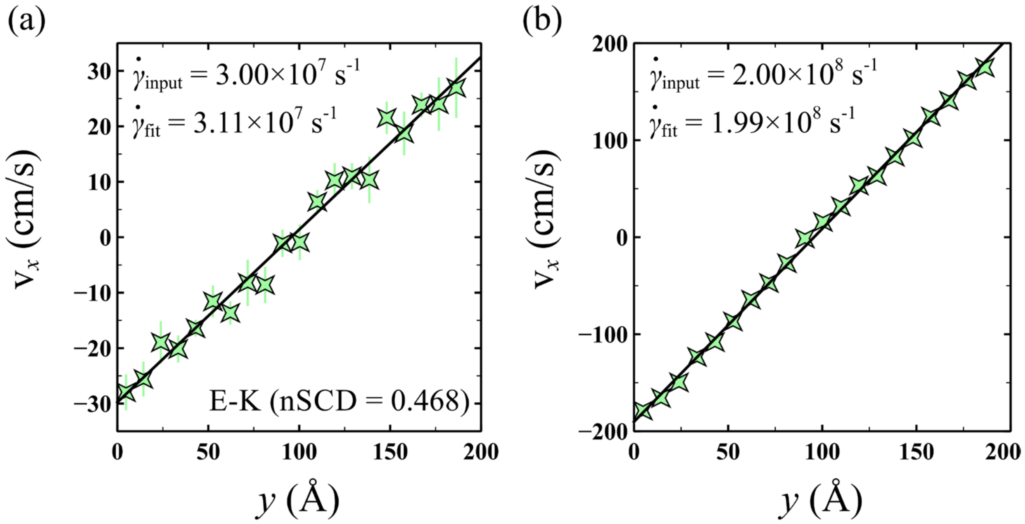


**Fig. S18.** Velocity $v_x$ of the residues in the flow direction $x$ as a function of gradient direction $y$ in the dense phase of a select E−K sequence with nSCD = 0.468 at (a) low and (b) high shear rates used in this work. The solid line corresponds to a linear fit to the data, the slope of which gives the shear rate $\dot{\gamma}_{\mathrm{fit}}$. The value of $\dot{\gamma}_{\mathrm{fit}}$ is nearly identical to the imposed shear rate $\dot{\gamma}_{\mathrm{input}}$ on the dense phase of the condensate.

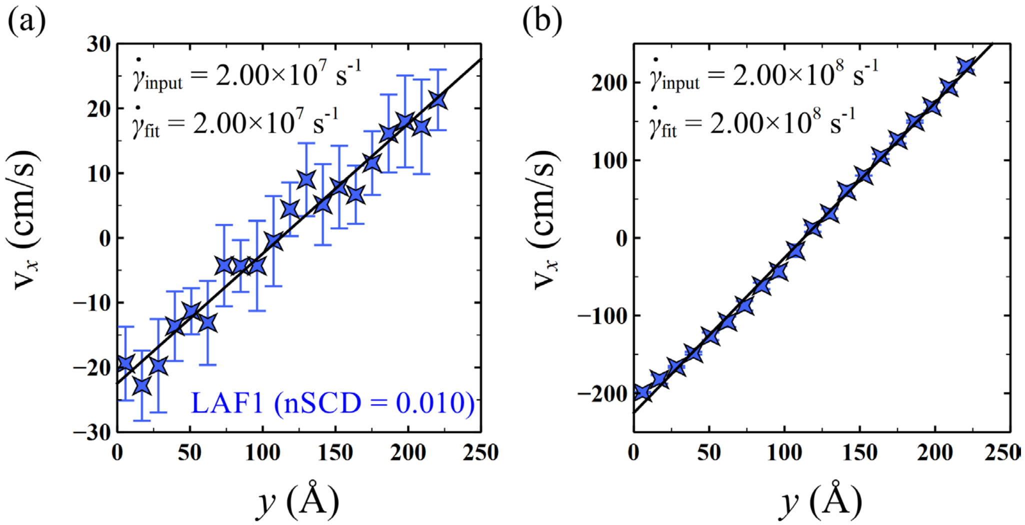


**Fig. S19.** Velocity $v_x$ of the residues in the flow direction $x$ as a function of gradient direction $y$ in the dense phase of a select LAF1 sequence with nSCD = 0.010 at (a) low and (b) high shear rates used in this work. The solid line corresponds to a linear fit to the data, the slope of which gives the shear rate $\dot{\gamma}_{\mathrm{fit}}$. The value of $\dot{\gamma}_{\mathrm{fit}}$ is nearly identical to the imposed shear rate $\dot{\gamma}_{\mathrm{input}}$ on the dense phase of the condensate.

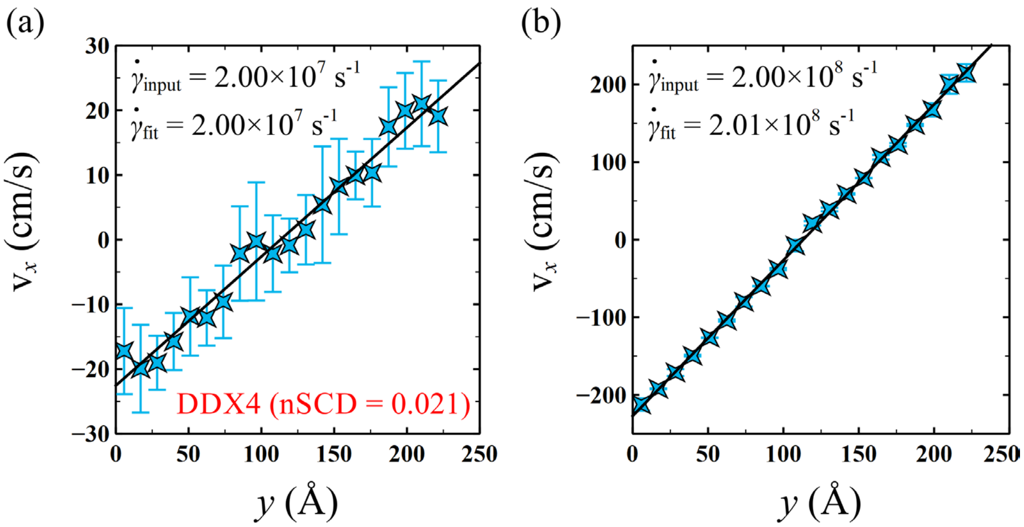


**Fig. S20.** Velocity $v_x$ of the residues in the flow direction $x$ as a function of gradient direction $y$ in the dense phase of a select DDX4 sequence with nSCD = 0.021 at (a) low and (b) high shear rates used in this work. The solid line corresponds to a linear fit to the data, the slope of which gives the shear rate $\dot{\gamma}_{\mathrm{fit}}$. The value of $\dot{\gamma}_{\mathrm{fit}}$ is nearly identical to the imposed shear rate $\dot{\gamma}_{\mathrm{input}}$ on the dense phase of the condensate.

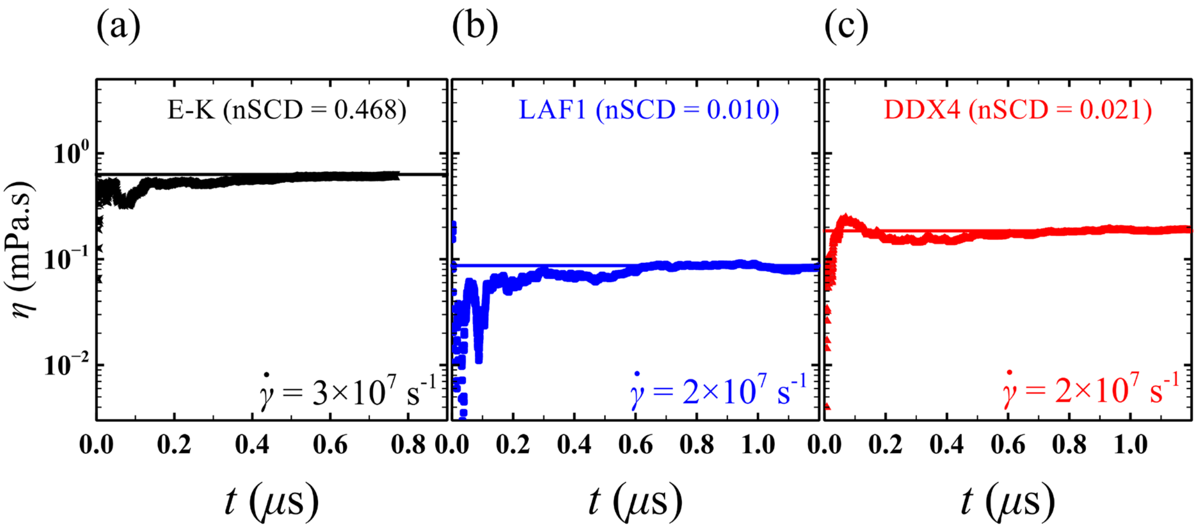


**Fig. S21.** Running average of shear viscosity $\eta$ as a function of simulation time $t$ at a low shear rate for select (a) E−K (nSCD = 0.468), (b) LAF1 (nSCD = 0.010), and (c) DDX4 (nSCD = 0.021) sequences. The solid lines represent the mean value of $\eta$ at the low shear rate for the respective sequences.

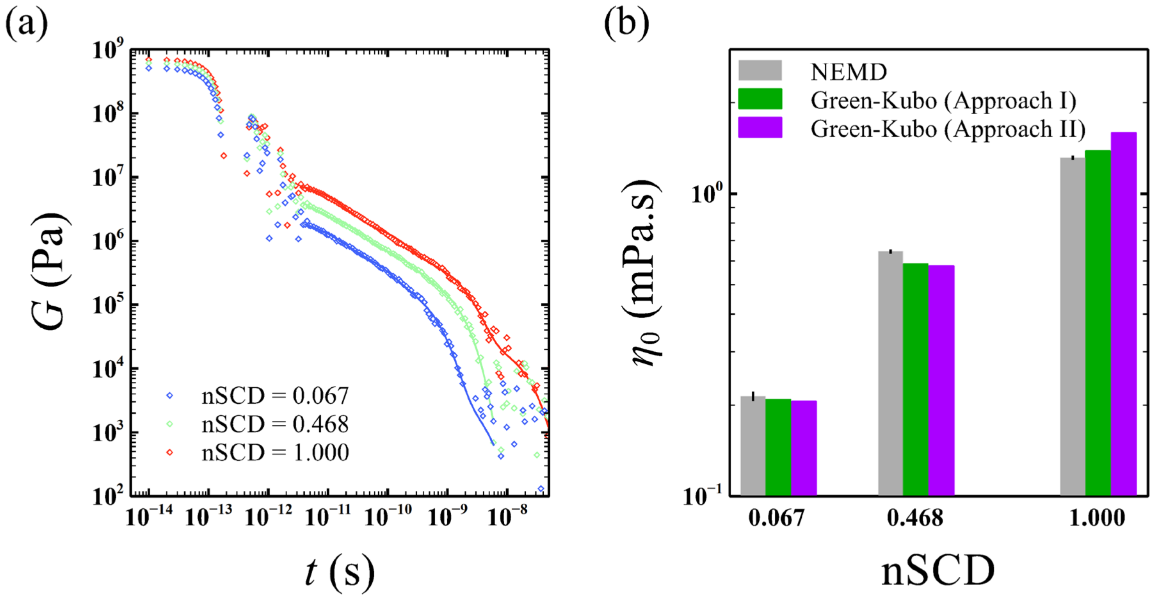


**Fig. S22.** (a) Shear stress relaxation modulus $G(t)$ for select E−K sequences. The solid lines at long time scales represent the fit to a series of Maxwell modes. (b) Comparison between zero-shear viscosity $\eta_0$ for the E−K sequences obtained from the NEMD simulations and the Green-Kubo relation. In approach I, $\eta_0$ was computed based on the Green-Kubo relation by simply integrating $G(t)$. In approach II, $\eta_0$ was computed as per the procedure outlined by Tejedor *et al.*[4]

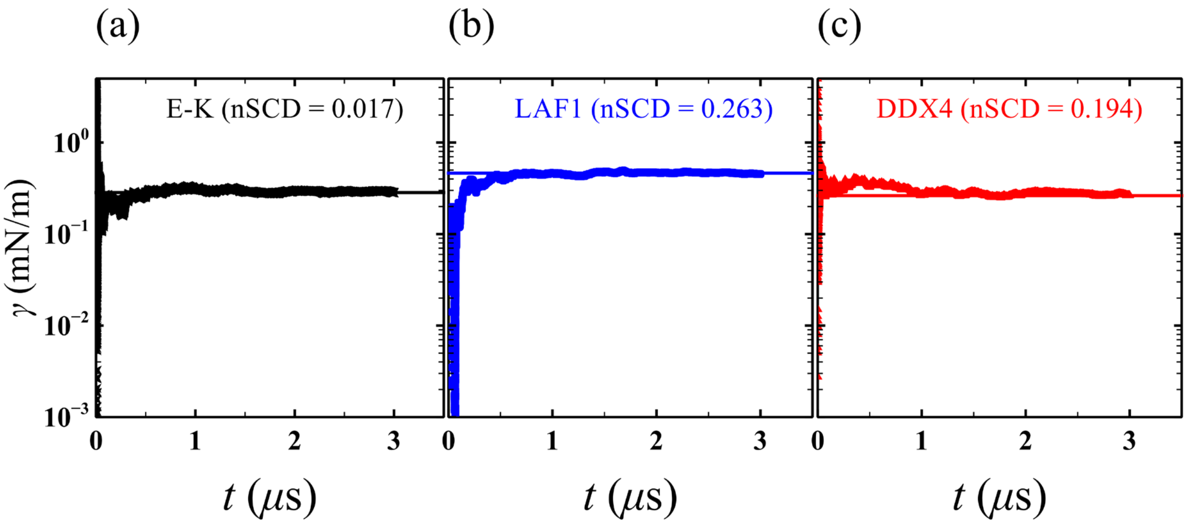


**Fig. S23.** Running average of surface tension $\gamma$ as a function of simulation time $t$ for select (a) E−K (nSCD = 0.017), (b) LAF1 (nSCD = 0.263), and (c) DDX4 (nSCD = 0.194) sequences. The solid lines represent the mean value of $\gamma$ for the respective sequences.

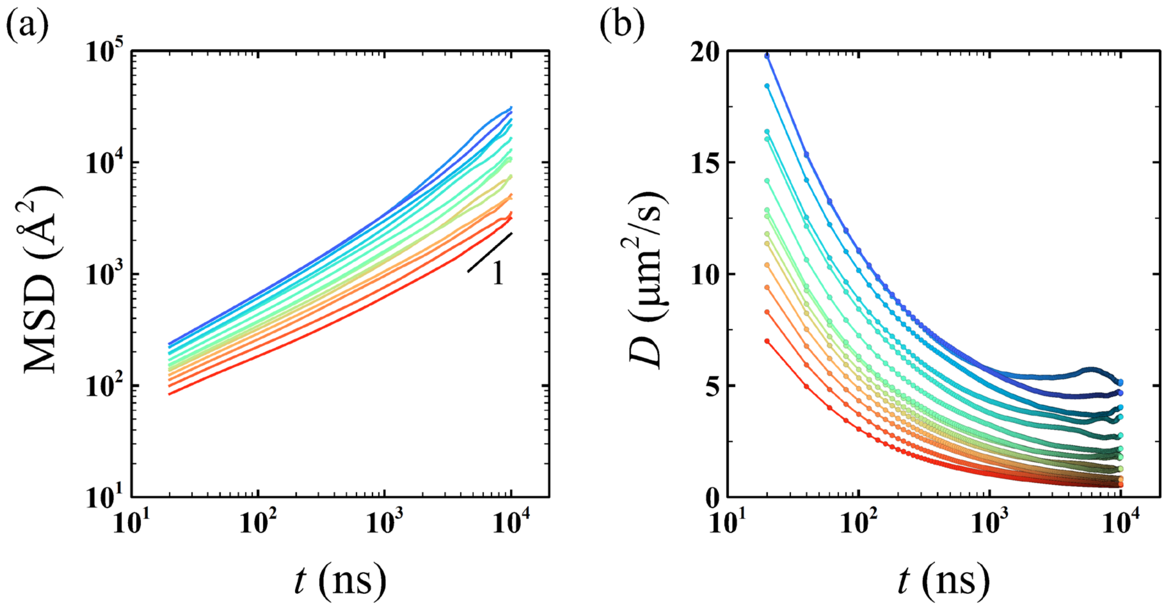


**Fig. S24.** (a) Mean square displacement MSD($t$) of the residues of a chain for the E−K sequences simulated using the Martini model. The slope value 1 indicates the diffusive regime. (b) Diffusion coefficient $D$ is extracted from the plateau region observed at long times ($t \geq 5$ $\mu$s) where the relation MSD $= 6Dt$ holds. The line color, ranging from blue to red, indicates increasing nSCD.

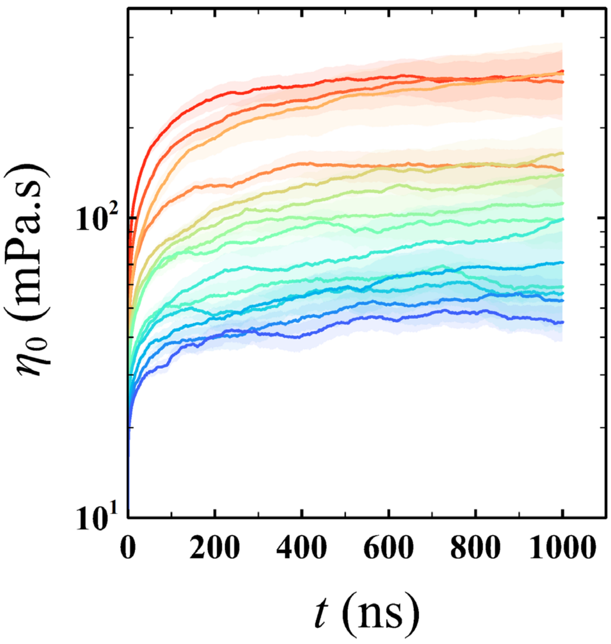


**Fig. S25.** (a) Zero-shear viscosity $\eta_0$ is obtained from the plateau region of the Green-Kubo integral observed at long times ($t \geq 0.5$ $\mu$s) for the E−K sequences simulated using the Martini model. The line color, ranging from blue to red, indicates increasing nSCD.

| Disordered sequence | $D$ ($\mu$m$^2/s$) from experiments* | $D$ ($\mu$m$^2/s$) from simulations# |
|---|---|---|
| LAF1 wild-type | $0.025 \pm 0.009$ | $0.052 \pm 0.002$ |
| DDX4 wild-type | $0.400 \pm 0.100$ | $0.061 \pm 0.003$ |

**Table S2.** Comparison of dense-phase diffusion coefficients $D$ between simulations (obtained based on the timescales mapped at the single-chain level at infinite dilution) and FRAP experiments for the LAF1 and DDX4 wild-type sequences. In FRAP, the recovery time of the fluorescently labeled protein molecules upon bleaching is measured to estimate the translational diffusion within the condensates. *The experimental conditions pertaining to the $D$ values for LAF1 and DDX4 were $T =$ 289.15 K to 291.15 K and $T = 293.15$ K, both in 150 mM NaCl buffer. #The simulation conditions pertaining to the $D$ values for LAF1 and DDX4 were $T = 280$ K and $T = 300$ K with Debye screening length $\ell = 10$ Å, which corresponds to 100 mM salt concentration.

| Protein sequence | $D_{\mathrm{MD}}$ based on intrinsic MD timescale ($\mathrm{m}^2/\mathrm{s}$) | $D_{\mathrm{sim}}$ in simulation units ($\sqrt{\varepsilon\sigma^2/m}$ or $\sigma^2/\tau$) | $D_{\mathrm{exp}}$ ($\mathrm{m}^2/\mathrm{s}$) | New timescale $\tau$ (s) from $D_{\mathrm{sim}} = D_{\mathrm{exp}}$ |
|---|---|---|---|---|
| E−K | $3.897 \times 10^{-7}$ | 27.99 | $1.571 \times 10^{-10}$ | $1.782 \times 10^{-9}$ |
| LAF1 | $1.385 \times 10^{-7}$ | 9.07 | $0.539 \times 10^{-10}$ | $1.683 \times 10^{-9}$ |
| DDX4 | $0.982 \times 10^{-7}$ | 6.45 | $0.867 \times 10^{-10}$ | $0.744 \times 10^{-9}$ |

**Table S3.** We first computed the diffusion coefficient $D_{\mathrm{MD}}$ of a single-chain based on the intrinsic MD timescale ($\tau$ = 48.89 fs) for the E−K, LAF1, and DDX4 sequences (second column). Note that $T =$ 300 K was used for the E−K and DDX4 sequences, while $T = 280$ K was used for the LAF1 sequences. Next, we represented the diffusion coefficient in simulation units $D_{\mathrm{sim}}$ (third column) by dividing $D_{\mathrm{MD}}$ by $\sqrt{\varepsilon\sigma^2/m}$ (or $\sigma^2/\tau$) for the E−K, LAF1, and DDX4 sequences, respectively. Average mass $m$ of each sequence and length scale $\sigma = 10^{-10}$ m were used in computing the values of $\sqrt{\varepsilon\sigma^2/m}$. We also calculated the experimentally expected diffusion coefficient $D_{\mathrm{exp}}$ of a single-chain in water as per the Stokes-Einstein relation (fourth column). Finally, we matched $D_{\mathrm{sim}}$ with $D_{\mathrm{exp}}$ to obtain new timescales $\tau$ (fifth column) that would yield the same single-chain diffusion coefficient in both simulations and experiments for the E−K, LAF1, and DDX4 sequences.

| Protein condensate | Protein molecular structure | Viscosity (Pa.s) | Surface tension (mN/m) | Brief method details to get surface tension |
|---|---|---|---|---|
| LAF1 RGG[3] | Fully disordered | $1.62 \pm 0.18$ Micropipette aspiration | $0.159 \pm 0.010$ Micropipette aspiration | From the intercept of the measured shear rate at different aspiration pressure |
| polyK[5] | Fully disordered | 0.204 SPT | 0.017 Fusion | Independent measure of viscosity is used in the surface tension to viscosity ratio obtained based on condensates fusion timescale |
| polyR[5] | Fully disordered | 14.4 SPT | 0.1 Fusion | Same as polyK |
| $[RGRGG]_5$ + dT40[7] | Fully disordered | 3 SPT | 0.8 Fusion | Same as polyK |
| polyK (pK) + heparin (H)[1] | Fully disordered | $0.30 \pm 0.03$ Oscillatory microrheology | $0.0571 \pm 0.0038$ Dual optical traps | Dual optical traps to stretch the condensates and the resulting static spring constant is proportional to surface tension |
| Proline-rich motif (P) + heparin (H)[1] | Fully disordered | $0.53 \pm 0.04$ Oscillatory microrheology | $0.067 \pm 0.0085$ Dual optical traps | Same as pK + H |
| PGL3[8] | Disordered + folded | 1 Dual optical traps | $\sim 4.5 \times 10^{-3}$ Dual optical traps | Dual optical traps to periodically stretch the condensates. Dynamic spring constant and complex shear modulus gives surface tension |
| FUS[9] | Disordered + folded | 0.7 Dual optical traps | $\sim 3.1 \times 10^{-3}$ Dual optical traps | Same as PGL3 |
| SH3 domain (S) + proline-rich motif (P)[1] | S: disordered + folded<br><br>P: fully disordered | $3.75 \pm 0.14$ Oscillatory microrheology | $0.0734 \pm 0.0058$ Dual optical traps | Same as pK + H |

| Protein condensate | Protein molecular structure | Viscosity (Pa.s) | Surface tension (mN/m) | Brief method details to get surface tension |
|---|---|---|---|---|
| SH3 domain (S) + lysozyme (L)[1] | S: disordered + folded<br><br>P: fully disordered | $10.1 \pm 1.1$<br>Oscillatory microrheology | $0.106 \pm 0.015$<br>Dual optical traps | Same as pK + H |
| NPM1[6] | Disordered + folded | 0.74<br>SPT | $8 \times 10^{-4}$<br>Fusion and sessile drop | Fusion: Same as polyK<br><br>Sessile drop: Condensates of various sizes were imaged through prism. Density, length scales from droplet shape, and gravity were used to obtain surface tension |

**Table S4.** Surface tension and viscosity of protein condensates and the experimental methods used to measure them in the literature. SPT corresponds to single particle tracking in which the mean square displacement of the fluorescent tracer beads embedded in the condensates is related to the condensate viscosity through the Stokes-Einstein relation.

| Protein condensate[1] | Fusion time (ms) | Viscosity (Pa.s) | Surface tension (mN/m) | Surface tension from viscocapillary model (mN/m) |
|---|---|---|---|---|
| polyK (pK) + heparin (H) | $30.3 \pm 0.9$ | $0.30 \pm 0.03$ | $0.0571 \pm 0.0038$ | 0.0585 |
| Proline-rich motif (P) + heparin (H) | $39.3 \pm 3.9$ | $0.53 \pm 0.04$ | $0.067 \pm 0.0085$ | 0.0797 |
| SH3 domain (S) + proline-rich motif (P) | $384 \pm 31$ | $3.75 \pm 0.14$ | $0.0734 \pm 0.0058$ | 0.0577 |
| SH3 domain (S) + lysozyme (L) | $1774 \pm 128$ | $10.1 \pm 1.1$ | $0.106 \pm 0.015$ | 0.0337 |

**Table S5.** Comparison of estimated surface tension based on the viscocapillary model with the directly measured values for four protein condensates formed by oppositely charged binary mixtures.
